# Method for Depletion of Mitochondria DNA in Human Bronchial Epithelial Cells

**DOI:** 10.1101/2023.07.28.551015

**Authors:** Michael V. Novotny, Weiling Xu, Anny Mulya, Allison J. Janocha, Serpil C. Erzurum

## Abstract

**Introduction:** Mitochondria are increasingly recognized to play a role in the airway inflammation of asthma. Model systems to study the role of mitochondrial gene expression in bronchial epithelium are lacking. Here, we create custom bronchial epithelial cell lines derived from primary airway epithelium that are depleted of mitochondrial DNA.

**Methods:** We treated BET-1A and BEAS-2B cells with ethidium bromide (EtBr) with or without 2’,3’-dideoxycytidine (ddC) to create cells lacking mitochondrial DNA (mtDNA). Cells’ mtDNA copy number were verified by quantitative polymerase chain reaction (qPCR) in comparison to nuclear DNA (nDNA). Cells were also assessed for oxidative phosphorylation by measures of oxygen consumption using the Seahorse analyzer.

**Results:** One week of EtBr treatment led to ∼95% reduction of mtDNA copy number (mtDNA-CN) in cells (mtDNA-CN, mean±SE, baseline vs. treatment: BEAS-2B, 820 ± 62 vs. 56 ± 9; BET-1A, 957 ± 52 vs. 73 ± 2), which was further reduced by addition of 25 μM ddC (mtDNA-CN: BEAS-2B, 2.8; BET-1A, 47.9). Treatment for up to three weeks with EtBr and ddC led to near complete loss of mtDNA (mtDNA-CN: BEAS-2B, 0.1; BET-1A, 0.3). The basal oxygen consumption rate (OCR) of mtDNA-depleted BET-1A and BEAS-2B cells dropped to near zero. Glycolysis measured by extracellular acidification rate (ECAR) increased ∼two-fold in cells when mtDNA was eliminated [ECAR (mpH/min/10^3^ cells), baseline vs. treatment: BEAS-2B, 0.50 ± 0.03 vs. 0.94 ± 0.10 *P*=0.005; BET-1A, 0.80 ± 0.04 vs. 1.14 ± 0.06 *P*=0.001].

**Conclusion:** Mitochondrial DNA–depleted BET-1A ρ0 and BEAS-2B ρ0 cell lines are viable, lack the capacity for aerobic respiration, and increase glycolysis. This cell model system can be used to further test mitochondrial mechanisms of inflammation in bronchial epithelial cells.

## Introduction

Mitochondria have their own circular DNA (mtDNA) that encodes 37 genes (13 proteins, 22 tRNAs, and 2 rRNAs) (1). Each human cell contains hundreds to thousands of mitochondria depending on tissue and cell type, and each mitochondrion has multiple copies of mtDNA (2). Variation of mtDNA copy number (mtDNA-CN), which is associated with mitochondrial enzyme activity and ATP production, can be used as an indirect biomarker of mitochondrial function (3, 4). mtDNA encodes 13 proteins of the electron transport chain; however, the majority of mitochondrial proteins are encoded by nuclear DNA (nDNA). Mitochondria independently replicate, transcribe, and translate their mtDNA using their own ribosomal and transfer RNA, which are encoded by mtDNA. A decrease in mtDNA can result in low levels of mitochondria-encoded proteins for oxidative phosphorylation and a mismatch of the levels of mtDNA-encoded proteins and nDNA-encoded proteins (3, 4).

Decrease in mtDNA-CN is associated with cardiovascular disease, diabetes, and cancers (3, 4). Mitochondrial changes have been mechanistically linked to asthma (5, 6). Previously, we reported that asthmatic airway epithelium has greater expression of nDNA-encoded mitochondrial respiratory chain complexes and hyper-dense supercoiled mitochondria (5). While the mtDNA-CN in platelets in a small group of asthmatics was not different than controls (7), others reported that mtDNA-CN of whole blood DNA was significantly higher in asthma as compared to controls in African Americans (8). While variation of mtDNA-CN in blood has been used in population studies, there are limited systems for studying the effect of loss of mtDNA in a cell-specific manner, such as in the airway epithelium.

Due to multiple copies of mtDNA, lack of a protective histone protein, and limited mtDNA repair capacity, mtDNA is more susceptible to excess reactive oxygen species (ROS) than nDNA. The non-coding regulatory D-loop region (mtDNA control region) is known to be most sensitive to ROS-mediated damage. Increased mtDNA D-loop methylation induced by excess ROS is associated with reduced mtDNA-CN and ultimately the deletion of the mtDNA (9).

Others have created cell lines without functional mtDNA (ρ0 cells). Currently, there are several available ρ0 cells, e.g., HeLa ρ0 (10), but these ρ0 cell lines were generated from immortal cancer cells. Creating ρ0 cells lines from non-tumorigenic sources will enable understanding of mitochondria in nonmalignant diseases. Development of primary airway cell line models will enable investigation of mitochondria, metabolism, and bioenergetics.

Here, we describe detailed methods for deletion of mtDNA from BET-1A and BEAS-2B bronchial epithelial cells, human cell lines derived from bronchial epithelial sources.

## Materials and Methods

### Cell Culture Conditions for BET-1A and BEAS-2B

BET-1A and BEAS-2B cells, a kind gift from Curtis Harris (NCI) (11), were grown in LHC-9+ 1% penicillin G (Pen) and streptomycin (Strep) (serum-free) medium (Gibco) following established protocols under sterile conditions (12). Cells were plated on tissue culture plates coated with a coating medium (1 ml Vitogen 100, 10 ml 10xBSA, 1 ml Fibronectin, and 100 ml LHC Basal medium). Cells were passaged with trypsin/EDTA at ratio 1:10. Cells are sensitive to traces of typsin/EDTA (13). Several precautions were taken by neutralizing trypsin with TNS (Trypsin Neutralizer Solution) and refreshing the culture medium 2 hours post plating.

### Cell Culture Conditions for HeLa and A549

HeLa cells were grown in Dulbecco’s Modified Eagle Medium (DMEM) + 10% fetal bovine serum (FBS) + 1% pen/strep medium and sub-cultured every 3-4 days. A549 cell were grown in Minimum Essential Medium (MEM) + 10% FBS + 1% pen/strep + 1% glutamine medium and sub-cultured every 3-4 days.

### Generation of ρ0 cells

To create mtDNA-free cells, cultured cells were treated with the mutagenic compound ethidium bromide (EtBr) alone or with addition of a nucleoside reverse transcriptase inhibitor ± 2’,3’-dideoxycytidine (ddC). The mutagenic compound is non-specific in its function, but nuclear DNA has a greater ability to repair itself compared to mitochondrial DNA (5). Reverse transcriptase inhibitors (like ddC) have been shown to specifically inhibit mitochondrial DNA (14). The detailed workflow is shown in Figure 1.

**Figure 1.**
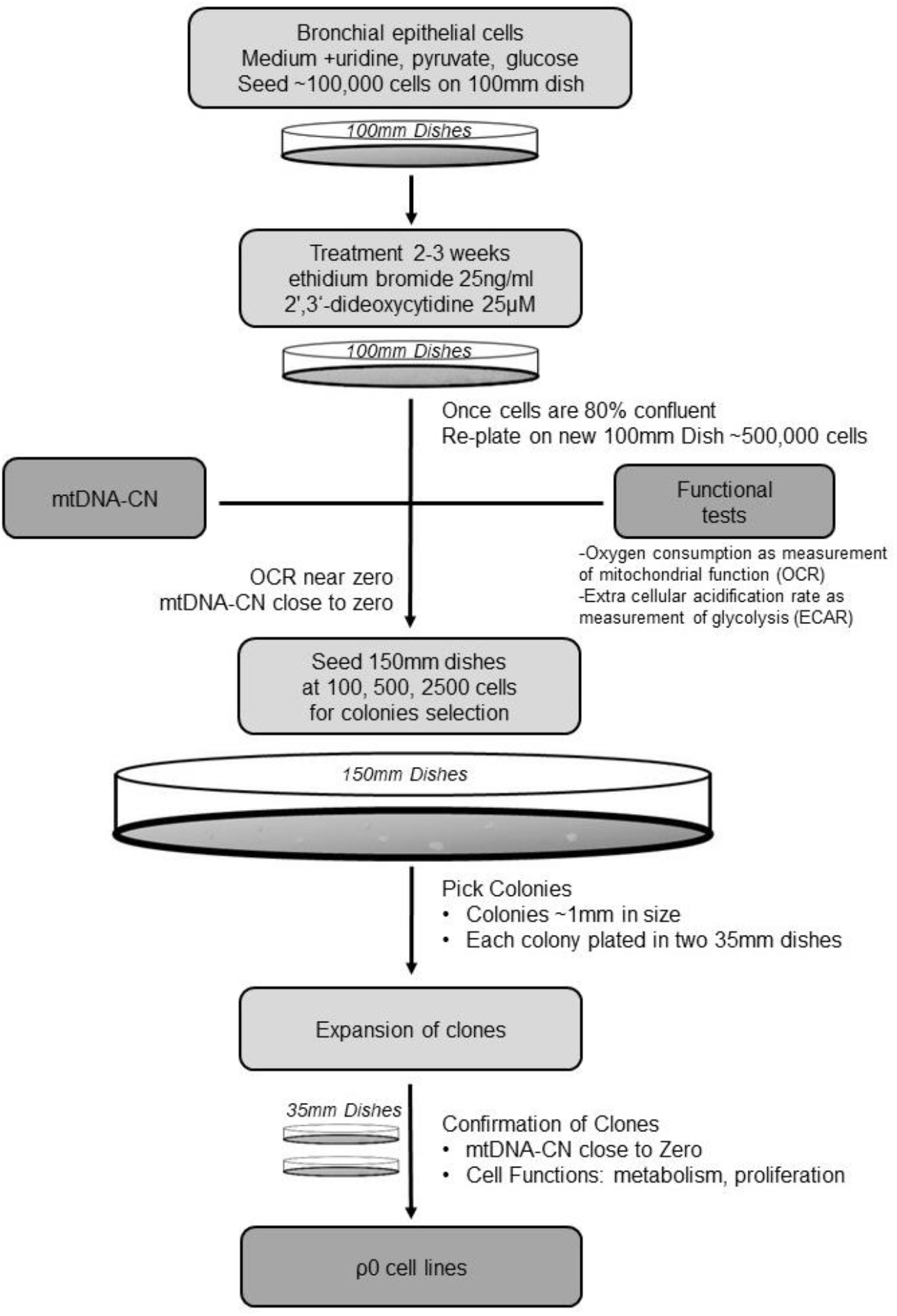
Methodology of creating ρ0 cells from BET-1A and BEAS-2B cells

BET-1A and BEAS-2B cells were initially plated at 100,000 cells on 100 mm tissue culture dishes in LHC-9 supplemented with 50 mg/ml uridine, 55 mg/ml pyruvate, and 3 mg/ml glucose. Cells were exposed to 25 or 50 ng/ml EtBr and 0, 25, or 50 ng/ml ddC (15). Media with EtBr and ddC was changed every other day. Cells were split when the cells became 80% confluent, and 500,000 cells were then plated on a new 100 mm tissue culture dish. The remaining cells were used to assess mtDNA-CN. Cells were grown under the given treatment conditions and tested until mtDNA-CN reached nearzero. At that point, cells were switched to media supplemented with uridine, pyruvate, and glucose without EtBr or ddC. The potential ρ0 cells were then split into 150 mm dishes at 100, 500, or 1000 cells per plate for single colony selection/isolation to derive a homogenous cell population. Clones were picked from plates and isolated in two 35 mm dishes for expansion. mtDNA-CN of expanded clones were verified by quantitative real-time Polymerase Chain Reaction (qPCR) and by gel electrophoresis, cellular respiration by Seahorse, and morphology via microscopy. Cells that persistently had near zero mtDNA-CN for two or more months of culture were designated BET-1A ρ0 or BEAS-2B ρ0

### DNA Extraction Cells and Tissues

Genomic DNA was extracted with the Qiagen DNeasy Blood & Tissue kit according to the manufacturer’s protocol [Qiagen, Hiden, Germany]. DNA amounts and concentrations were determined using Invitrogen Qubit 3 system with Qubit dsDNA Broad Range kit (Thermo Fisher Scientific, Waltham, MA USA). DNA from human lung and heart samples were obtained from explanted tissues during transplantations under Cleveland Clinic IRB1546 with informed consent.

### Determination of Optimal Primer Targets for mtDNA and nDNA

We used five sets of previously reported primers that target either nuclear or mtDNA genes (Table 2); multiple annealing primer pair sets were purchased from Integrated DNA Technologies (IDT) [Coralville, Iowa USA] for mitochondrial / nuclear DNA targets. A mitochondrial DNA monitoring primer set kit from Takara Bio [San Jose, CA USA] was purchased for comparison. Dried oligos were hydrated with water to 100 nM concentration. In a new 1.5ml tube, forward and reverse primer sets were mixed at a 1:1 ratio to make a 10x stock for qPCR reactions.

**Table 1.**
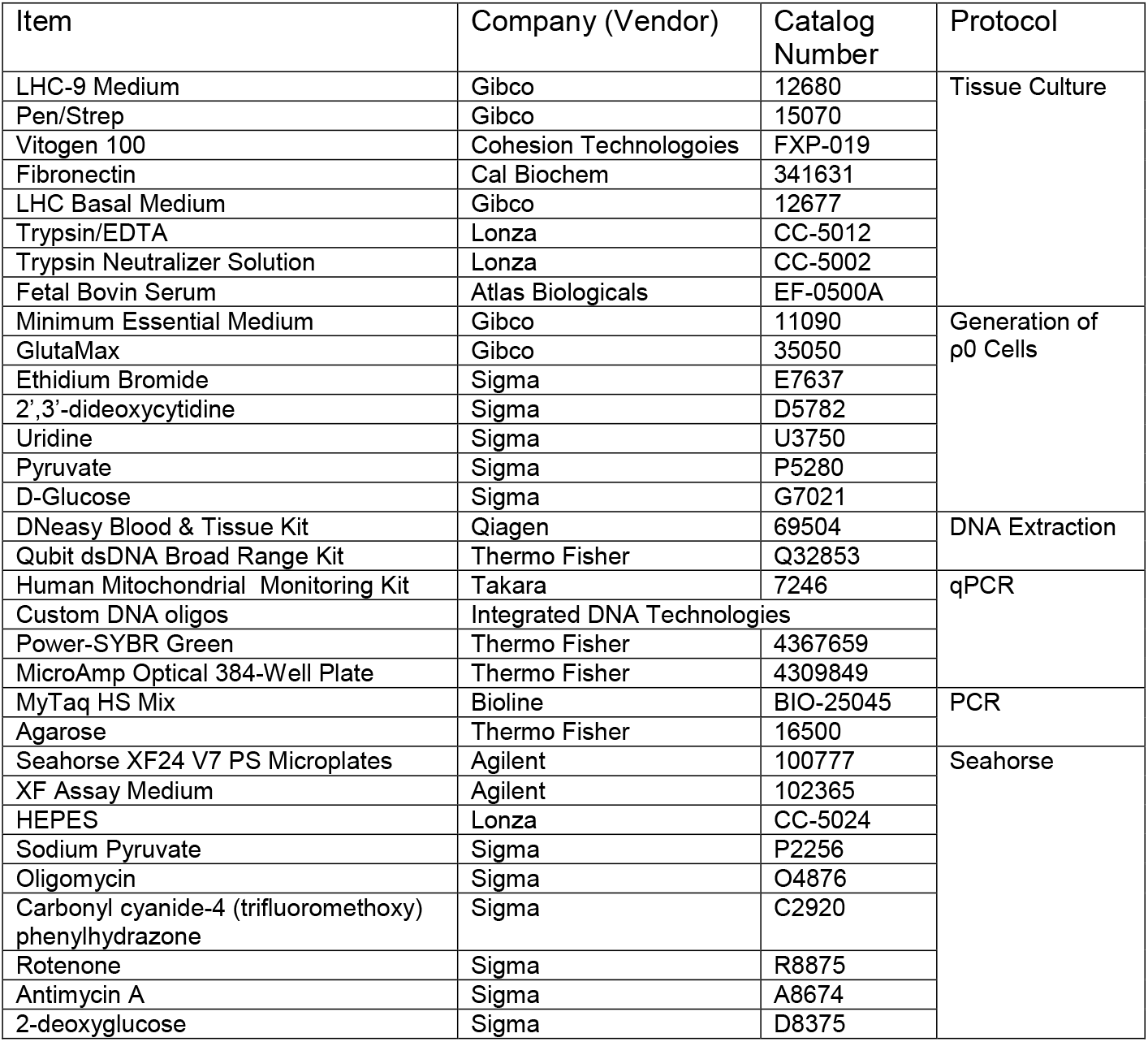
List of reagents, vendors, and catalog numbers.

**Table 2.**
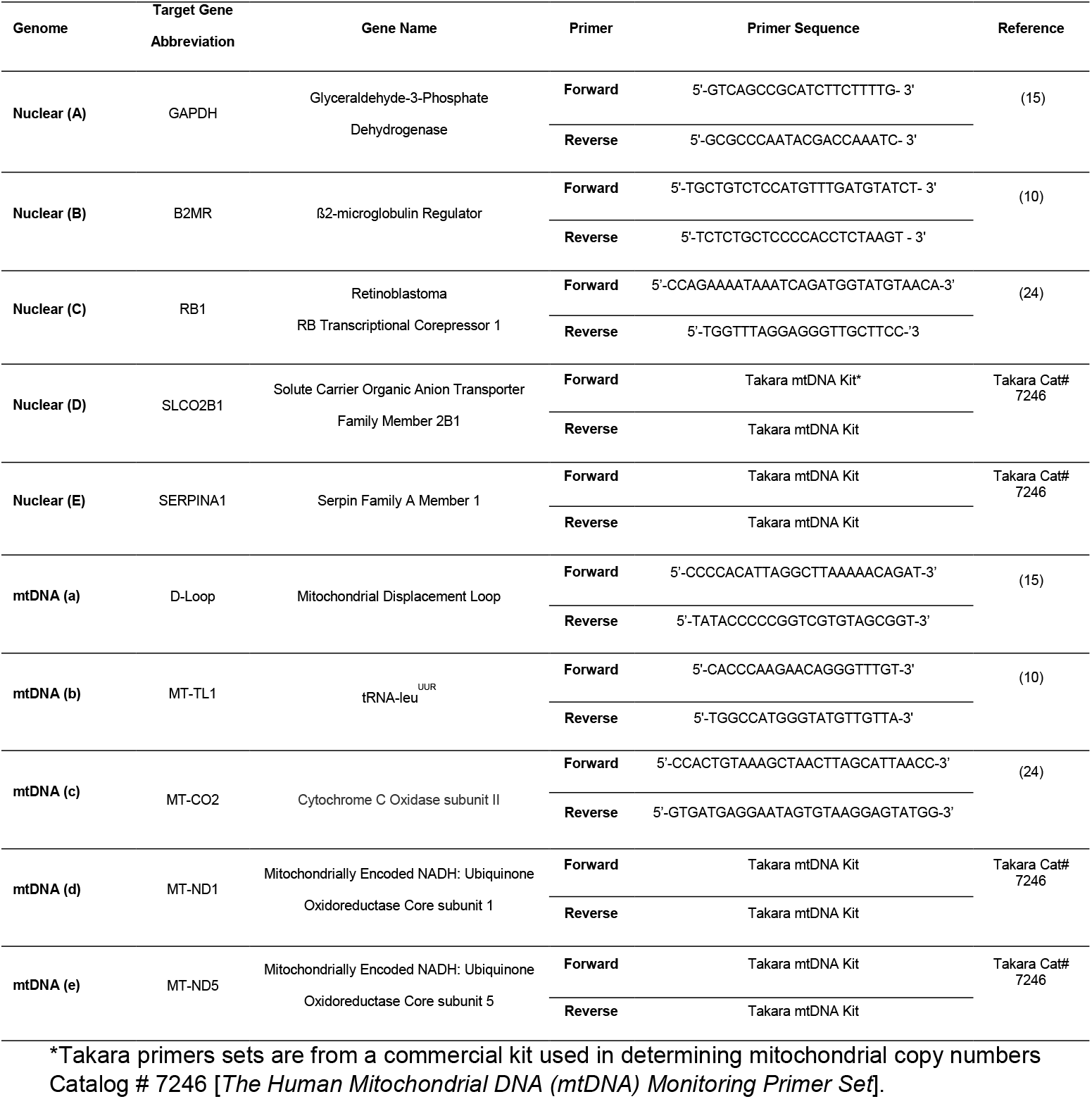
Primer sequences for target genes.

### Monitoring Mitochondrial DNA Elimination by qPCR

ρ0 cells have mtDNA-CN close to zero (0.1 is typically reported in HeLa ρ0) (10). We aimed to have a similarly low number in our BET-1A and BEAS-2B ρ0 cells.

The qPCR reaction was performed in triplicate for each sample at a volume of 20 μl per well. Each qPCR reaction contained 2 μl diluted sample DNA (varying concentrations of DNA tested as described below), 10 μl Power-SYBR Green Polymerase Chain Reaction (PCR) Master Mix (Applied Biosystems), 6 μl PCR water, and 2 μl primer mix containing 10 nM forward and reverse primers. qPCR reactions were performed on Applied Biosystems QuantStudio5 using MicroAmp Optical 384-Well Reaction Plates. The realtime PCR conditions were the following: initial denaturation 2 minutes at 50°C ramp to 95°C for 10 minutes followed by 40 amplification cycles of (95°C for 15 seconds denaturation ramp down to 60°C for 60 secs for annealing and extension) followed by a final 95°C for 15 second hold. The cycle threshold values (Ct values) were determined automatically via QuantStudio Design & Analysis software.

To optimize the qPCR methods, we tested several primer sets detecting both mtDNA and nDNA. Primer sets tested are reported in Table 1. For this purpose, master mix was prepared by mixing DNA, SYBR-Green and PCR water, and then added to the PCR plates which contain the 2 μL of primer sets.

### Quantification of mtDNA Copy Number Using qPCR

mtDNA copy number was quantified in relation to nDNA. The average Ct values of qPCR targeting mtDNA were subtracted from the average Ct values of qPCR targeting the nDNA; this was defined as ΔCt. mtDNA-CN is calculated with the following formula: 2 (each PCR cycle represents doubling the amount) to the power of the ΔCt value and multiply by the number of homologous chromosomes (2 in normal primary cells).

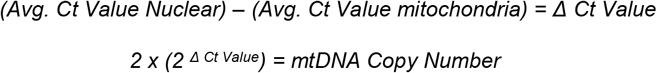

### Absence of mtDNA by Gel Electrophoresis

To confirm the loss of mtDNA, DNA from BET-1A ρ0 and BEAS-2B ρ0 cells were compared to the original non-treated source cell line (BET-1A and BEAS-2B) by PCR and gel electrophoresis. PCR reactions were performed in a volume of 25 μl per well on a XYX Technik A6 Thermal Cycler PCR machine with mtDNA tRNA-leu (MT-TL1) and nuclear DNA GAPDH primer sets. Each PCR reaction contained 2 μl annealing primer set, 12.5 μl MyTaq HS mix (Meridian Bioscience, Memphis, TN), 2 μl DNA template (5 ng total), and 8.5 μl PCR water. The PCR conditions were as follows: initial denaturation to 94°C for 1 minute followed by 25 amplification cycles of (94°C for 30 seconds denaturation, ramp down to 55°C 30 secs for annealing, and 72°C for 30 seconds extension) followed by a final 72°C for 5 minutes ramp down to 4°C hold. A 1.5% ultrapure agarose gel in 1X TAE buffer was cast. 10 μl PCR sample and 5 μl 1kb DNA ladder were loaded in each well. Gels were run for 45 mins at 80 volts, and the image was documented with the Bio-RAD ChemiDoc MP Imaging System (Bio-Rad, Hercules, CA USA)

### Measurement of Oxygen Consumption Rate (OCR) and Extracellular Acidification Rate (ECAR) in Cells

When qPCR copy numbers indicated ρ0 sample, loss of electron transport chain activity was evaluated with a functional assay to measure cellular respiration and energy metabolism (bioenergetics).

Assessment of cellular bioenergetics was performed using the extracellular flux analyzer XFe24 (Seahorse Biosciences, Agilent, Billerica, MA) with a modified MitoStress Test protocol. BET-1A, BEAS-2B, BET-1A-ρ0, and BEAS-2B-ρ0 cells were plated on a precoated Seahorse XF24 V7 PS cell culture microplate with 30×10^3^ cells per well. Cells were cultured in the medium with supplements (uridine, pyruvate, and glucose) overnight. On the day of the assay, growth medium was removed, cells were washed with XF assay medium, and the media was replaced with XF assay medium. XF assay medium was serum-free DMEM supplemented with 5 mM HEPES, 5.6 mM D-glucose, 2 mM L-glutamine, and 1 mM sodium pyruvate in the absence of sodium bicarbonate, and pH of the media was adjusted to 7.4. Oxygen consumption rate (OCR) and extracellular acidification rate (ECAR) were measured at baseline, in response to 1 μM oligomycin (port A), 1 μM carbonyl cyanide-4 (trifluoromethoxy) phenylhydrazone (FCCP) (port B), 1 μM rotenone and antimycin A (port C), and 100 mM 2-deoxyglucose (port D) for three measurements each. After extracellular flux analysis, cells were stained with 1.6 μM of Hoechst 33342 stain (Thermo Fisher Scientific, Waltham, MA USA), and cell numbers were determined with the Cytation 5 (Biotek Instruments, Agilent, Winooski, VT USA). OCR and ECAR data were expressed as pmol/minute/10^3^ cells and mpH/minute/10^3^ cells, respectively. Data are mean ± SEM of at least 5 replicates.

## Results

### Validity of mtDNA Copy Number Quantification

Primer sets for mtDNA and nDNA targets for routine testing were established. There is expected variability in Ct value amounts for housekeeping nDNA genes when comparing to mtDNA genes. As the number of primer sets used increases, accuracy increases, but the quantity and cost of each PCR reaction also increases. qPCR was run with five different nuclear and mitochondrial targets (Figure 2). Pair set combinations were compared with the average of all five annealing sets for both nuclear and mitochondrial targets to determine which pair would be ideal for mtDNA-CN rigor and reproducibility. Based on Ct values in Figure 2, the following primer pairs were chosen for mtDNA [tRNA-Leu^UUR^ (MT-TL1) and Cytochrome C Oxidase subunit II (MT-COII)]. For nDNA, GAPDH and ß2-microglobulin (Glob) were chosen.

**Figure 2.**
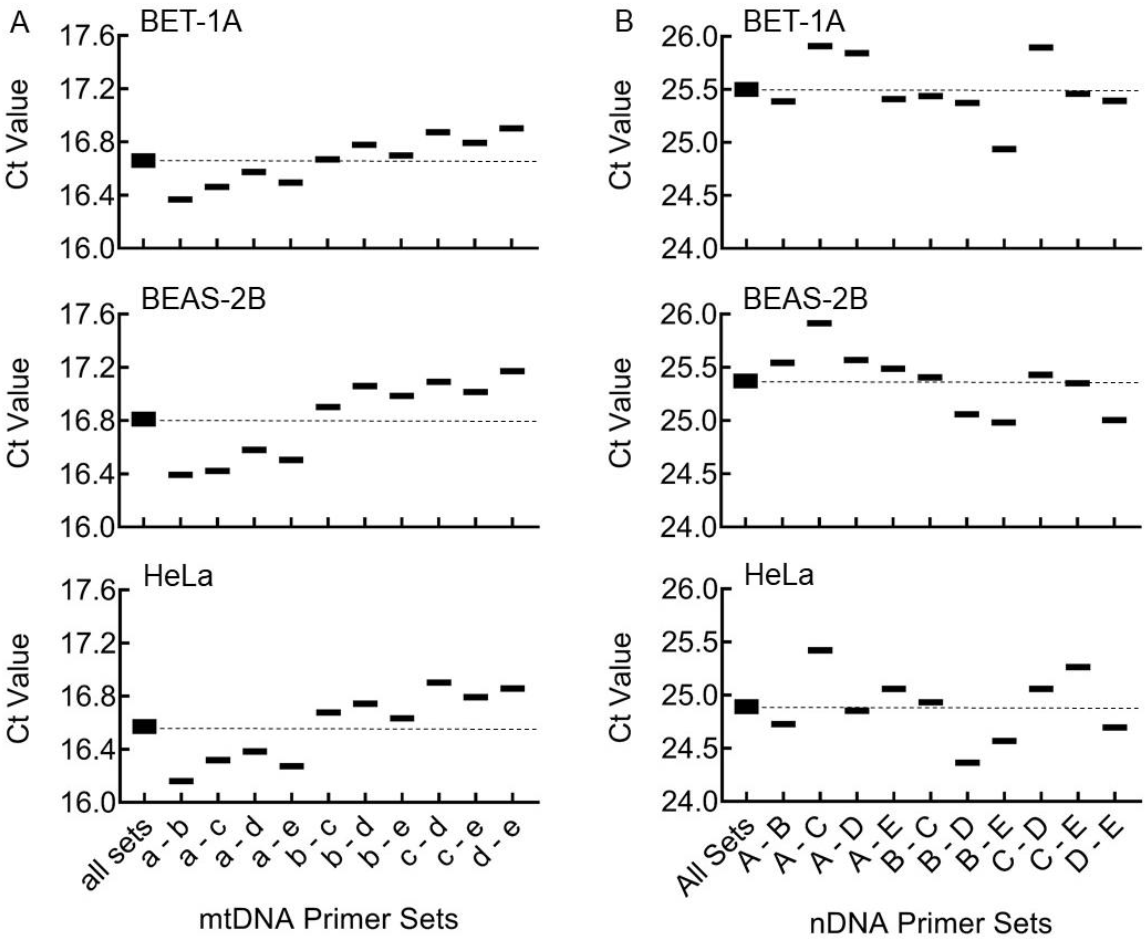
Determining primers pairs for mtDNA-CN calculation. Cycle thresholds for various combination of primers were used to determined the ideal pair to represent the sample. (A) Average Ct values of five sets of mitochondrial primers (all sets) were compared to various combinations of two sets of Ct values on BET-1A, BEAS-2B, or HeLa mtDNA. (B) Average Ct values of five sets of nuclear primers (All Sets) were compared to various combinations of two sets of Ct values on BET-1A, BEAS-2B, or HeLa nDNA. A pair of primers were chosen as a representative value of the five sets, allowing future PCR to be performed with two pair sets for each nuclear or mitochondrial Ct value determination. Dotted line represents average Ct values of the five sets. See Table 1 for primer reference.

We also assessed whether variability of the DNA template amount would affect the calculated mtDNA-CN. DNA from six different source samples (BET-1A, BEAS-2B, HeLa, human lung tissue, human heart tissue, and the A549 cell line) were extracted and logarithmically diluted [0.1, 0.3, 1, 3, 10, 30, 100 ng] for mtDNA-CN determination (Figure 3). The calculated mtDNA-CN were consistent within each sample type from 0.1 to 30 ng DNA input. This indicates that any variation in DNA template that might occur within 0.1 – 30 ng will not significantly alter our mtDNA-CN calculations.

**Figure 3.**
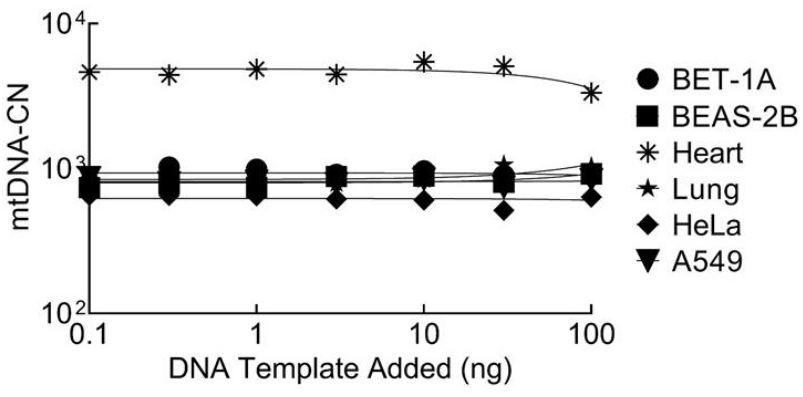
mtDNA-CN with various amount of DNA template. BET-1A, BEAS-2B, human heart/lung tissue, HeLa and A549 DNA were logarithmically diluted [0.1, 0.3, 1, 3, 10, 30, 100 ng] and used as a template for qPCR to determine mtDNA copy numbers.

### Generating ρ0 Cells

The process for creating ρ0 BET-1A and BEAS-2B cells required treating the cells with both EtBr (a mutagen) and ddC (a nucleoside reverse transcriptase inhibitor). Two to three weeks of EtBr treatment led to ∼95% reduction of mtDNA copy numbers in both cell lines, which was further reduced by the addition of at least 25 uM ddC (Figure 4 A/B). While the loss of 95% mtDNA copy numbers is significant, creating long-term ρ0 cells requires near complete loss of mtDNA. This only occurred in cells that were treated with both EtBr and ddC, i.e., mtDNA-CN reached <1 copy per nDNA. EtBr treated cells did not maintain low mtDNA-CN when treatment was discontinued. Once a near loss of mtDNA-CN was achieved, cells were plated for single colonies and allowed to expand with continued testing for mtDNA-CN. To confirm cells lacked mtDNA by a method other thanqPCR, DNA was amplified via PCR for mtDNA and nDNA targets and evaluated for PCR products by agarose electrophoresis (Figure 4 C/D). The band targeted for mitochondrial (MT-TL1) was absent in the newly created BET-1A ρ0 and BEAS-2B ρ0, while band targeted for nDNA (GAPDH) was still intact in BET-1A ρ0 and BEAS-2B ρ0. Established HeLa and HeLa ρ0 cells were used as controls.

**Figure 4.**
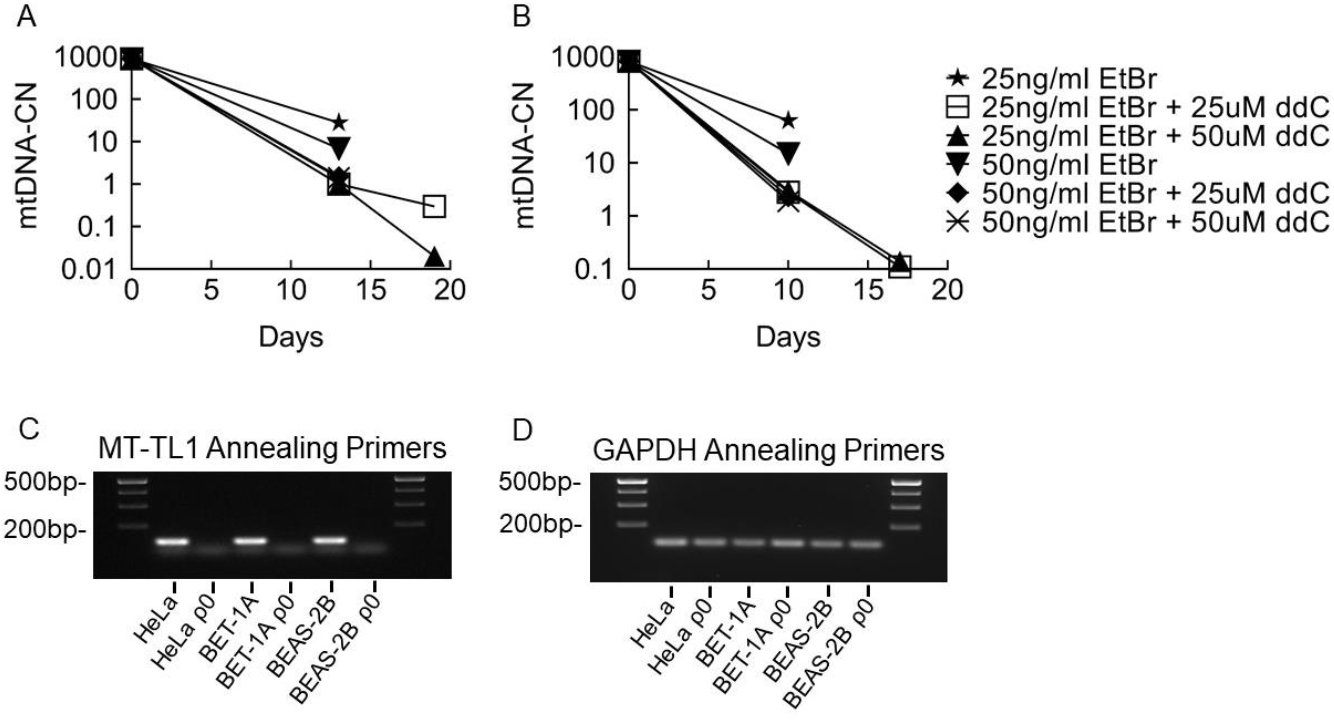
Depletion of mtDNA-CN in BET-1A and BEAS-2B ρ0. Analysis of mtDNA-CN over a 2-3 week period with different treatment conditions. BET-1A (A) and BEAS-2B (B) cells were treated with either 25 or 50 ng/ml EtBr +/-25 or 50 uM ddC on day 0. This graph tracks the progress of mtDNA copy numbers over a 2-week period under different treatment conditions. Some isolated cells were pelleted, and DNA was extracted, which in turn was used as the template for qPCR to determine mtDNA copy numbers. Amplification of MT-TL1 and GAPDH in cells detected by PCR. 1.5% agarose gel from PCR using tRNA-leu^UUR^ (C) and GAPDH (D) annealing primers.

### Validation of ρ0 Cell Lines

To confirm loss of functional electron transport chain, we performed Seahorse extracellular flux analysis. BET-1A and BEAS-2B have OCR and ECAR profiles typical of airway cells (16). However, the basal OCR values of mtDNA-depleted BET-1A and BEAS-2B cells were basically undetectable [OCR (pmol/min/10^3^ cells), baseline vs. treatment: BEAS-2B, 4.68 ± 0.12 vs. 0.06 ± 0.07 *P*<0.001 and N=4 (baseline) 7(treatments), technical replicates; BET-1A, 6.55 ± 0.30 vs. 0.02 ± 0.10 *P*<0.001 and N=4 (baseline), 7 (treatments), technical replicates]. ECAR values increased significantly in ρ0 cells [ECAR (mpH/min/1000 cells), baseline vs. treatment: BEAS-2B, 0.50 ± 0.03 vs. 0.94 ± 0.10 *P*=0.005 and N=4 (baseline), 7 (treatments), technical replicates; BET-1A, 0.80 ± 0.04 vs. 1.14 ± 0.06 *P*=0.001 and N=4 (baseline), 7 (treatments), technical replicates]. Interestingly, injection of oligomycin (the ATP synthase inhibitor) did not increase the ECAR in BET-1A or BEAS-2B ρ0 cells (Fig 5 A-B). These suggest that the cells were shifting energy metabolism from oxidative phosphorylation to glycolytic metabolism. These new ρ0 cells were multiplying and maintaining their shape and size typical of regular BET-1A cells (Fig 5 C-D). Visibly, the cells in culture were proliferating and had no significant difference in viability.

**Figure 5.**
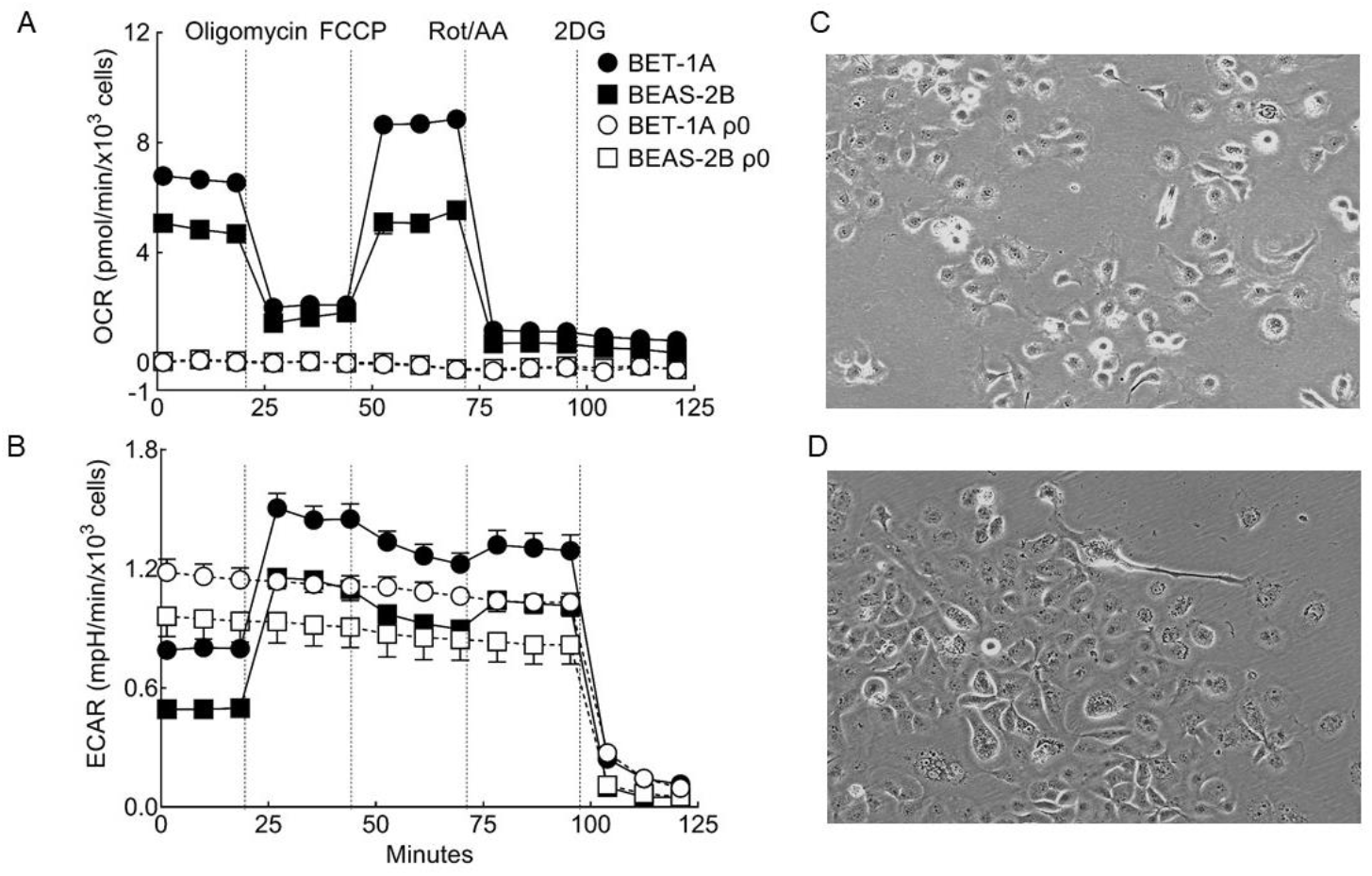
Glycolysis replaces oxidative phosphorylation as the source of energy in BET-1A p0 and BEAS-2B p0 cells. Assessment of cellular energy metabolism was performed using the XFe24 extracellular flux analyzer (Seahorse). OCR (A) and ECAR (B) profiles are shown for BET-1A (dark circle), BEAS-2B (dark square), BET-1A p0 (open circle), and BEAS-2B p0 (open square) at baseline, in response to oligomycin, FCCP, rotenone/antimycin A, and 2-deoxyglucose (2DG). Data are mean ± SEM of at least three replicates. BET-1A (C) and BET-1A ρ0 (D) cells in culture (100x magnification).

## Discussion

Creating non-tumorigenic derived cell lines without mitochondrial DNA can be accomplished with common research tools and techniques, opening up the potential of generating novel ρ0 cell lines. qPCR measurement of mtDNA copy numbers is a quick and reliable method to determine loss of mitochondria, while quantitative measures of oxygen consumption can validate the mtDNA deletion. Furthermore, the BET-1A and BEAS-2B cells depleted of mtDNA are viable and utilize greater glycolysis for energy production.

Before creating ρ0 cells, establishing a protocol for determining mitochondrial copy numbers requires choosing annealing primer sets targeting mitochondrial DNA and nuclear housekeeping genes. While there will be variable Ct value results depending on the target choice, as long as those gene targets are constant, comparative sample analyses can be accomplished. There are several factors to consider when selecting which genes to target. For nuclear primer targets, standard single chromosome housekeeping genes that remain constant under multiple conditions are ideal (17) (18) (19). Having a housekeeping gene not affected by treatments will allow different sample sets to be compared. For mitochondrial targets, we determined that looking at two targets would be prudent as opposed to a single mtDNA target. By testing multiple mtDNA targets, we confirm that the entire mtDNA is eliminated. When creating primer sets for both nuclear and mtDNA, melting temperatures of all primer sets should be similar (∼60°C). For our mtDNA targets, the average Ct values of MT-TL1 and MT-COII Ct values were consistently close to the average of five Ct values tested in three cell lines (BET-1A, BEAS-2B, and HeLa) and was the prime criteria for selecting mtDNA primers. For our nuclear targets, the average Ct values of GAPDH and ß2-microglobulin were used based on these being well-established stable housekeeping genes.

We optimized the minimum amount of DNA needed for mtDNA determination in order to retain the greatest number of cells for ongoing culture. When we performed a dilution curve on our DNA as a template, there was little difference in our calculated mitochondrial copy number in the six different sample types within the range of 0.1 to 30 ng DNA template. This provides confidence that despite any range of starting DNA, findings would still be accurate.

A limitation of SYBR-Green qPCR is less reliability and sensitivity as compared to TaqMan (20) (21). To address this, we used diluted DNA standards (HeLa and HeLa ρ0) to assess reproducibility across multiple qPCR runs (data not shown). The TaqMan chemistry system has a greater sensitivity and more consistent results, but the downside of TaqMan is greater cost of annealing primer sets. Thus, for qPCR to determine copy numbers in this methodology requiring multiple testing, SYBR-Green was a cost-effective method for determining loss of mtDNA. Unfortunately, methodology plays a critical role in quantitating mtDNA copy numbers, and different annealing targets and variable chemistry make it difficult to compare mtDNA copy numbers in cells across various reports. Nevertheless, the average numbers of mtDNA in BET-1A and BEAS-2B is ∼800 mtDNA to nDNA gene, which is comparable to prior reports in cancer cell lines of ∼1000 mtDNA to 1 nDNA (22). All data shown in this manuscript were preformed with SYBR-Green chemistry.

This report shows that it is possible to effectively delete mitochondrial DNA from transformed bronchial epithelial cells without compromising cell viability. In our first attempt to create BET-1A and BEAS-2B ρ0, the treatment condition was only using EtBr to mutate the mtDNA. This did decrease mtDNA-CN by ∼95% in two to three weeks.

However once EtBr treatment was discountinued, cells recovered mtDNA to original levels (data not shown). In the BET-1A and BEAS-2B cells, we determined that 25 ng/ml EtBr with 25 μM ddC was ideal to generate ρ0 cells. This resulted in cells with near-zero mtDNA and oxygen consumption. Considering the nature of EtBr to cause lasting damage to nuclear DNA and since the results of both tested concentrations were similar, using a lower EtBr dose combined with ddC is prudent to avoid nuclear damage (23). Once the treatment phase of creating ρ0 cell was successful, plating for single colonies on larger dishes at various amounts led to stable clones of cells. Overall, this method is highly successful for creation of ρ0 bronchial epithelial cells. These novel cell lines will be important in future studies to determine the role of mitochondria in bronchial epithelial functions.

## Acknowledgments

We are thankful to Suzy A. A. Comhair for providing needed human tissue samples, Lori Mavrakis for providing tissue culture expertise, and to Curtis Harris for the BEAS-2B and BET-1A. This work is supported in part by HL060917.

## References

1. Boore JL. Animal mitochondrial genomes. Nucleic Acids Res. 1999;27(8):1767–80.

2. Anderson S, Bankier AT, Barrell BG, de Bruijn MH, Coulson AR, Drouin J, et al. Sequence and organization of the human mitochondrial genome. Nature. 1981;290(5806):457–65.

3. Wallace DC. A mitochondrial bioenergetic etiology of disease. J Clin Invest. 2013;123(4):1405–12.

4. Castellani CA, Longchamps RJ, Sun J, Guallar E, Arking DE. Thinking outside the nucleus: Mitochondrial DNA copy number in health and disease. Mitochondrion. 2020;53:214–23.

5. Xu W, Ghosh S, Comhair SA, Asosingh K, Janocha AJ, Mavrakis DA, et al. Increased mitochondrial arginine metabolism supports bioenergetics in asthma. The Journal of clinical investigation. 2016;126(7):2465–81.

6. Reddy PH. Mitochondrial Dysfunction and Oxidative Stress in Asthma: Implications for Mitochondria-Targeted Antioxidant Therapeutics. Pharmaceuticals (Basel). 2011;4(3):429–56.

7. Xu W, Cardenes N, Corey C, Erzurum SC, Shiva S. Platelets from Asthmatic Individuals Show Less Reliance on Glycolysis. PLoS One. 2015;10(7):e0132007.

8. Cocco MP, White E, Xiao S, Hu D, Mak A, Sleiman P, et al. Asthma and its relationship to mitochondrial copy number: Results from the Asthma Translational Genomics Collaborative (ATGC) of the Trans-Omics for Precision Medicine (TOPMed) program. PLoS One. 2020;15(11):e0242364.

9. Kaarniranta K, Pawlowska E, Szczepanska J, Jablkowska A, Blasiak J. Role of Mitochondrial DNA Damage in ROS-Mediated Pathogenesis of Age-Related Macular Degeneration (AMD). Int J Mol Sci. 2019;20(10).

10. Khozhukhar N, Spadafora D, Rodriguez Y, Alexeyev M. Elimination of Mitochondrial DNA from Mammalian Cells. Curr Protoc Cell Biol. 2018;78(1):20 11 1–20 11 4.

11. Reddel RR, Ke Y, Gerwin BI, McMenamin MG, Lechner JF, Su RT, et al. Transformation of human bronchial epithelial cells by infection with SV40 or adenovirus-12 SV40 hybrid virus, or transfection via strontium phosphate coprecipitation with a plasmid containing SV40 early region genes. Cancer research. 1988;48(7):1904–9.

12. Uetani K, Arroliga ME, Erzurum SC. Double-stranded rna dependence of nitric oxide synthase 2 expression in human bronchial epithelial cell lines BET-1A and BEAS-2B. Am J Respir Cell Mol Biol. 2001;24(6):720–6.

13. Zhao F, Klimecki WT. Culture conditions profoundly impact phenotype in BEAS-2B, a human pulmonary epithelial model. J Appl Toxicol. 2015;35(8):945–51.

14. Nelson I, Hanna MG, Wood NW, Harding AE. Depletion of mitochondrial DNA by ddC in untransformed human cell lines. Somat Cell Mol Genet. 1997;23(4):287–90.

15. Schubert S, Heller S, Loffler B, Schafer I, Seibel M, Villani G, et al. Generation of Rho Zero Cells: Visualization and Quantification of the mtDNA Depletion Process. Int J Mol Sci. 2015;16(5):9850–65.

16. Xu W, Janocha AJ, Leahy RA, Klatte R, Dudzinski D, Mavrakis LA, et al. A novel method for pulmonary research: assessment of bioenergetic function at the air-liquid interface. Redox Biol. 2014;2:513–9.

17. Kim BR, Nam HY, Kim SU, Kim SI, Chang YJ. Normalization of reverse transcription quantitative-PCR with housekeeping genes in rice. Biotechnol Lett. 2003;25(21):1869–72.

18. Turabelidze A, Guo S, DiPietro LA. Importance of housekeeping gene selection for accurate reverse transcription-quantitative polymerase chain reaction in a wound healing model. Wound Repair Regen. 2010;18(5):460–6.

19. Dheda K, Huggett JF, Bustin SA, Johnson MA, Rook G, Zumla A. Validation of housekeeping genes for normalizing RNA expression in real-time PCR. Biotechniques. 2004;37(1):112-4, 6, 8-9.

20. Tajadini M, Panjehpour M, Javanmard SH. Comparison of SYBR Green and TaqMan methods in quantitative real-time polymerase chain reaction analysis of four adenosine receptor subtypes. Adv Biomed Res. 2014;3:85.

21. Cao H, Shockey JM. Comparison of TaqMan and SYBR Green qPCR methods for quantitative gene expression in tung tree tissues. J Agric Food Chem. 2012;60(50):12296–303.

22. Siegismund CS, Schafer I, Seibel P, Kuhl U, Schultheiss HP, Lassner D. Mitochondrial haplogroups and expression studies of commonly used human cell lines. Mitochondrion. 2016;30:236–47.

23. Nass MM. Differential effects of ethidium bromide on mitochondrial and nuclear DNA synthesis in vivo in cultured mammalian cells. Exp Cell Res. 1972;72(1):211–22.

24. Leibowitz RD. Effect of Ethidium Bromide on Mitochondrial DNA Synthesis and Mitochondrial DNA Structure in Hela Cells. J Cell Biol. 1971;51(1):116-+.

